## Supplemental materials for "The biochemical activities of the Saccharomyces cerevisiae Pif1 helicase are regulated by its N-terminal domain"

**The N-terminal domain of Saccharomyces cerevisiae Pif1 is necessary to regulate telomerase activity**

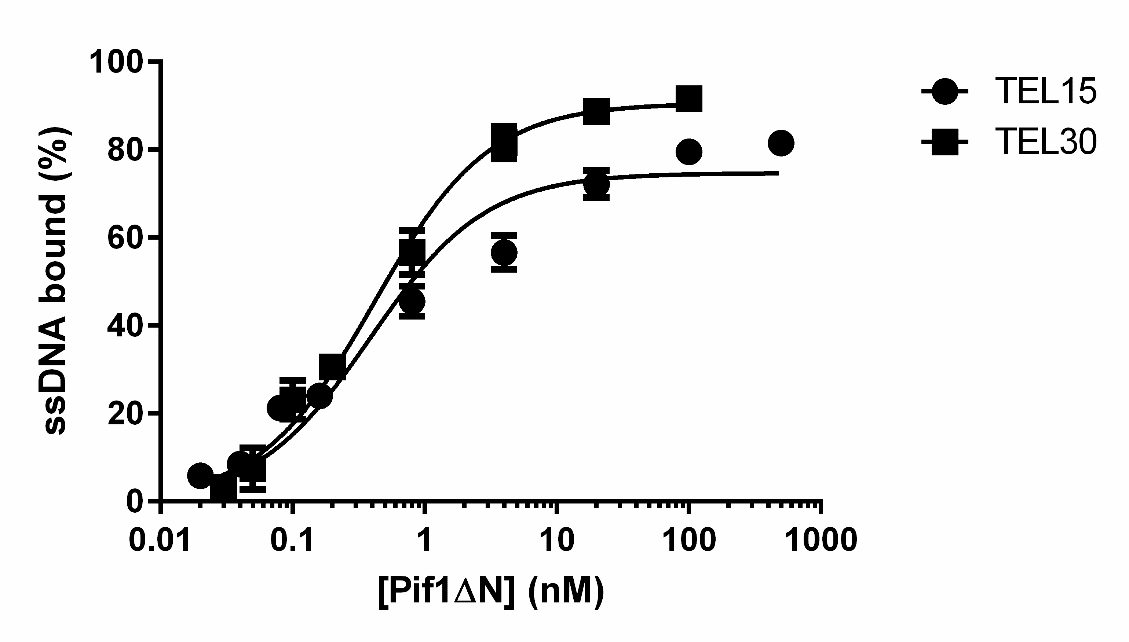

**A**

**B**

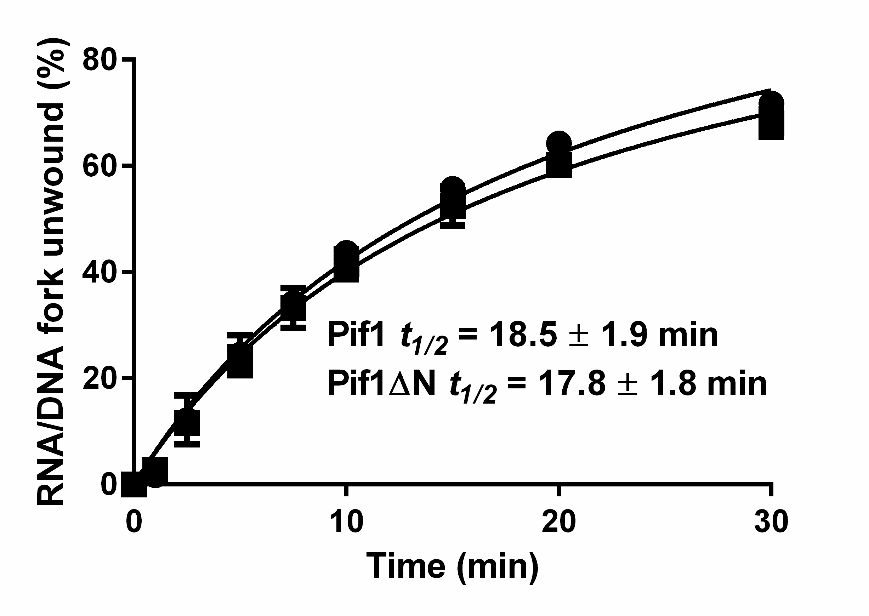

**Supplemental Figure S1. Pif1ΔN binds ssDNA tightly and unwinds DNA with similar kinetics to full-length Pif1. A**) Pif1ΔN binds the Tel15 and Tel30 ssDNA substrates with similar affinity. The plotted values represent the results of EMSAs using radiolabeled substrates and the indicated concentrations of recombinant Pif1ΔN protein. **B**) Pif1 and Pif1ΔN unwind forked DNA with similar kinetics. The plotted values represent DNA unwinding time course assays performed with 0.4 nM of a radiolabeled DNA-DNA fork substrate and 2 nM Pif1 or Pif1ΔN. In all cases, the data represent the averages of ≥ 3 independent experiments, and the error bars are the standard deviation.
